# A co-transcriptional ribosome assembly checkpoint controls nascent large ribosomal subunit maturation

**DOI:** 10.1101/2023.03.02.530842

**Authors:** Zahra A. Sanghai, Rafal Piwowarczyk, Arnaud Vanden Broeck, Sebastian Klinge

**Author notes:** Correspondence should be addressed to S.K.

## Abstract

During transcription of eukaryotic ribosomal DNA in the nucleolus, assembly checkpoints exist that guarantee the formation of stable precursors of small and large ribosomal subunits. While the formation of an early large subunit assembly checkpoint precedes the separation of small and large subunit maturation, its mechanism of action and function remain unknown. Here, we report the cryo-electron microscopy structure of the co-transcriptional large ribosomal subunit assembly intermediate that serves as a checkpoint. The structure provides the mechanistic basis for how quality control pathways are established through co-transcriptional ribosome assembly factors, that structurally interrogate, remodel, and together with ribosomal proteins cooperatively stabilize correctly folded pre-ribosomal RNA. Our findings thus provide a molecular explanation for quality control during eukaryotic ribosome assembly in the nucleolus.

---

Eukaryotic ribosome assembly involves the coordinated activity of more than 200 assembly factors that assist in the formation of small and large ribosomal subunits^1^. During the transcription of rDNA, a 35S precursor transcript is generated in which the small subunit rRNA (18S) and two of the three large subunit (LSU) rRNAs (5.8S and 25S) are flanked by external transcribed spacers (5’ ETS and 3’ETS) and interspersed by internal transcribed spacers (ITS1 and ITS2). A unique and poorly understood challenge during co-transcriptional stages of ribosome biogenesis in the nucleolus is the correct assembly and stabilization of the 5’ ends of nascent pre-ribosomes via co-transcriptional ribosome assembly factors.

Miller spreads have highlighted the co-transcriptional appearance of assembly intermediates as terminal structures that first form for the small subunit where the synthesis of a 5’ ETS ribonucleoprotein (RNP) precedes the assembly of the small subunit processome^2-4^. The co-transcriptional formation of an uncharacterized LSU assembly intermediate around the 5’ end of the LSU pre-ribosomal RNA (pre-rRNA) is then followed by pre-rRNA cleavage at site A2 within ITS1, which separates small and large subunit maturation^2^.

However, how the structural integrity of the 5’ segment of the nascent LSU precursor is interrogated has so far remained elusive. It is unclear how ribosome assembly factors implicated in this event can perform key checkpoint functions that require the presence and recognition of distinct pre-rRNA elements to facilitate the separation of small and large subunit biogenesis.

The Noc1-Noc2 complex together with Rrp5 has been implicated in mediating some of these functions as well as co-transcriptional cleavage at site A2, since Rrp5 is involved in both small and large subunit assembly^5-10^. Chemical biology approaches have highlighted that the Noc1-Noc2 complex and Rrp5 can stably bind to pre-ribosomal RNA mimics that contain a 5’ segment of the LSU^11,12^. However, the mode by which this complex may establish a checkpoint that can control the coordinated progression of both small and large subunit assembly has so far remained unclear^13^.

To characterize the mechanisms by which nascent LSUs are chaperoned in a co-transcriptional manner, we employed a system wherein pre-rRNA mimics are generated from a plasmid *in vivo*^3,12,14^. This system allowed for the synthesis of a pre-rRNA mimic starting at site A2 in ITS1 and terminating after domain VI of the 25S rRNA, followed by a set of MS2 RNA aptamers as previously described^12^ **(Extended Data Fig. 1a)**. The expression of this pre-rRNA mimic resulted in the formation of mature LSUs, showing that this pre-rRNA mimic undergoes the entire LSU assembly pathway (**Extended Data Fig. 1b,c**). By using either Noc1 or Noc2 as baits for affinity purification, we were able to isolate both an early co-transcriptional assembly intermediate containing the Noc1-Noc2 complex (hereafter referred to as the Noc1-Noc2 RNP) and a known late nucleolar pre-60S assembly intermediate (State E, containing Noc2-Noc3)^15^, indicating our pre-rRNA mimic proceeds along a physiologically relevant assembly pathway (**Extended Data Figs. 1, 2**). To determine the structure of the Noc1-Noc2 RNP, extensive 3D classification was performed on eight cryo-EM datasets to first obtain a well-defined module containing domain II of the 25S rRNA before resolving a more flexible domain I and 5.8S rRNA module **(Extended Data Figs. 1e-g, 3, 4, 5)**.

The resulting composite cryo-EM reconstruction contains domains I and II of the 25S rRNA, the 5.8S rRNA and ITS2. This arrangement of domains is expected for the earliest co-transcriptional ribosome assembly intermediate visualized in Miller spreads, as the protein composition of subsequent intermediates only changes significantly once domain VI has been transcribed and assembled for post-transcriptional maturation^11,12^ (**Fig. 1a, b**). The overall structure of this complex highlights that the domain I and II modules act as largely independent units that, in contrast to the subsequent assembly states, are not yet joined through ribosomal proteins and rRNA^15-17^. In agreement with prior biochemical data, the 5.8S rRNA is largely flexible with the most ordered regions corresponding to segments that base-pair with the 25S rRNA^18^.

**Figure 1.**
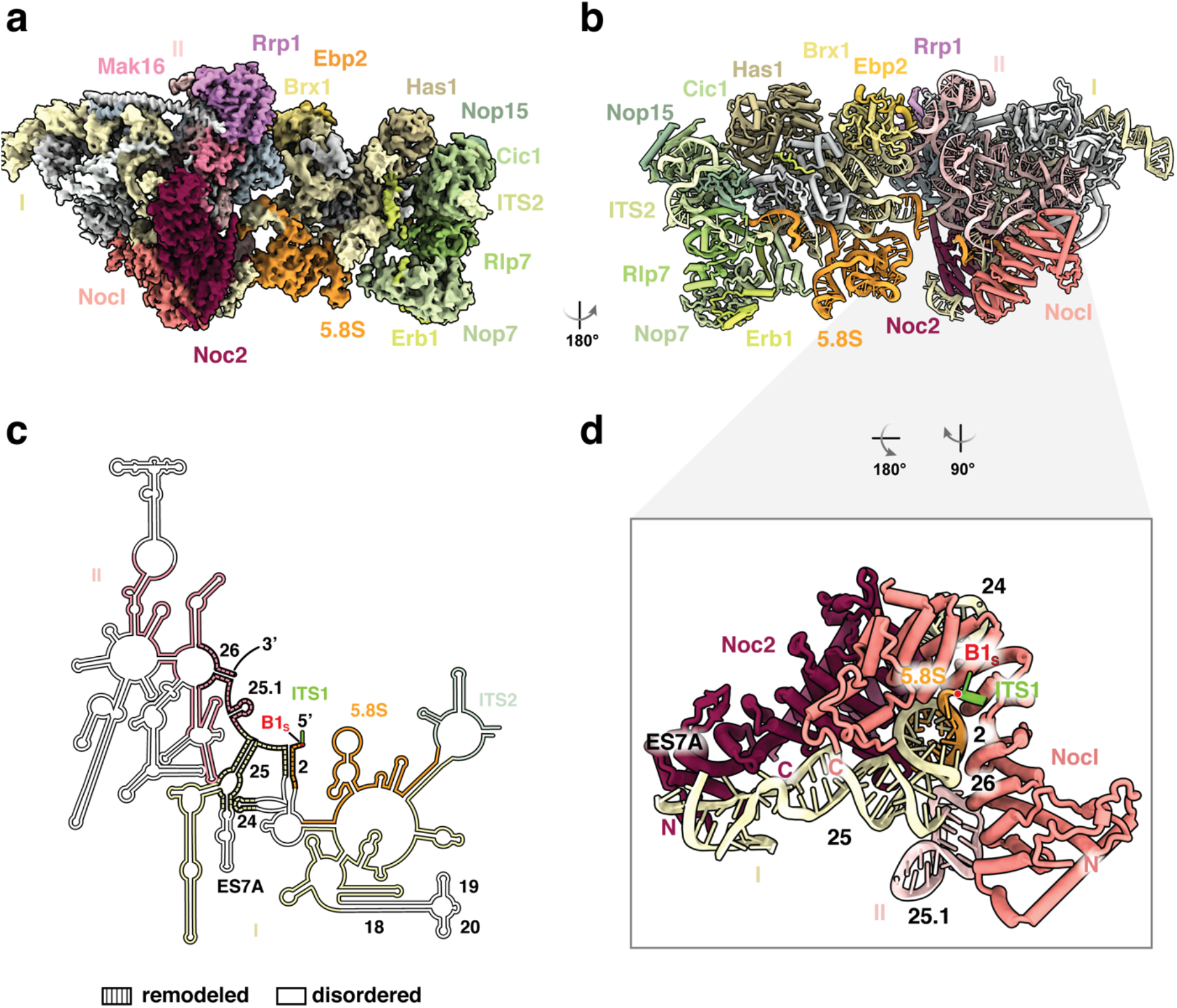
Cryo-EM structure of the co-transcriptional Noc1-Noc2 RNP. **(a)** Cryo-EM reconstruction of the Noc1-Noc2 RNP, visualizing domains I and II of the 25S rRNA with ITS2 and the 5.8S rRNA, and associated assembly factors and ribosomal proteins. **(b)** Corresponding atomic model of the Noc1-Noc2 RNP rotated by 180°. **(c)** rRNA secondary structure diagram of the yeast LSU rRNA visualized in the Noc1-Noc2 RNP cryo-EM reconstruction. rRNA elements present in the structure are color-coded, disordered segments colored in white, and remodeled segments indicated by dashed lines, with relevant helices numbered in black. **(d)** The Noc1-Noc2 dimer clamps around root helix 2 of the pre-rRNA formed by base-pairing of 5.8S and 25S rRNAs and remodels root helices 25 and 26.

The most prominent feature of this particle is the Noc1-Noc2 complex, which is responsible for remodeling all root helices formed by the 5.8S rRNA and domains I and II of the 25S rRNA (helices 2, 25 and 26) (**Fig. 1c, d, Extended Data Fig. 6**). Since root helices form the architectural core of the LSU^19,20^, their early recognition by the Noc1-Noc2 complex indicates a key point of quality control. The Noc1-Noc2 complex not only binds to several critical root helices, but specifically encircles helix 2 through molecular shape complementarity in a non-sequence specific manner (**Fig. 1d**). Helix 2 is a crucial root helix of the forming LSU: it contains the 5’ end of the LSU pre-rRNA and initiates the formation of the polypeptide exit tunnel, one of the essential functional centers of the LSU, which is formed in a tightly controlled manner during subsequent assembly stages in the nucleolus^15-17^. While the Noc1-Noc2 complex is evolutionarily related to the Noc2-Noc3 and Nop14-Noc4 complexes, it is the only member of this family that can directly chaperone pre-ribosomal RNA, in this case helix 2 of the nascent LSU pre-rRNA (**Extended Data Figs. 7, 8)**. The captured state further contains an unprocessed 5’ end of the LSU pre-rRNA as we observe a part of ITS1 that extends beyond the B1_S_ cleavage site, which later generates the mature 5’ end of the 5.8S rRNA (**Fig. 1d, Extended Data Fig. 6c**).

The strategic position of Noc1-Noc2 within the Noc1-Noc2 RNP is key to its primary function during the establishment of a co-transcriptional ribosome assembly checkpoint. In conjunction with assembly factors Mak16 and Rrp1, Noc1-Noc2 pre-assembles the highly intertwined rRNA domains I and II, thereby positioning these domains for co-operative stabilization by ribosomal proteins (**Fig. 2a, Extended Data Fig. 9**). Beyond the immediate formation of the co-transcriptional ribosome assembly checkpoint, the Noc1-Noc2 complex further controls post-transcriptional ribosome assembly at four levels: 1) RNA topology, 2) RNA processing, 3) the incorporation of ribosomal proteins, and 4) the chronology of ribosome assembly factor association (**Fig. 2**).

First, at the level of RNA topology, the Noc1-Noc2 complex not only separates domains I and II for independent maturation, but also prevents the integration of domain VI, which is a post-transcriptional event (**Figs. 1, 2, Extended Data Fig. 9**)^15-17^. Key links between domains I and II, such as helix 19 are prevented from establishing contacts and Noc1 substantially overlaps with domain VI, thereby preventing its incorporation during early co-transcriptional stages of large subunit assembly (**Fig. 2, Extended Data Fig. 9)**.

Second, given the position of Noc1 with respect to the 5’ end of the 5.8S rRNA, Noc1 may also serve as physical barrier preventing exonucleolytic trimming of ITS1 beyond site B1_S_ (**Fig. 1d, 2a**).

Third, at the level of ribosomal protein incorporation, the structure rationalizes previous genetic and biochemical data showing that RPL17 only gradually associates with early LSU assembly intermediates, as its binding site is initially occupied by Noc1^21^ (**Fig. 2, Extended Data Fig. 9c-f**). Similarly, the Noc2 C-terminus interferes with the incorporation of RPL26 as it prevents the ordering of helix 19 of the 25S rRNA, thereby eliminating a key binding site for RPL26 at the junction of domains I and II. The importance of critical interfaces between domains I, II and the 5.8S rRNA is further highlighted by the finding that key stabilizers of these sites include Diamond-Blackfan anemia proteins RPL17, RPL26 and RPL35A (**Extended Data Fig. 9c-f**)^22,23^.

Fourth, at the level of ribosome assembly factor chronology, the Noc1-Noc2 dimer is presumably recruited to early LSU assembly intermediates via an interaction with the Rrp5 N-terminus, which could not be assigned in our reconstruction. While Noc2 contacts Mak16, it prevents the Mak16-associated ribosome assembly factor complex Rpf1-Nsa1 from binding near domains I and VI (**Fig. 2**). In addition, Noc1 sterically clashes with the Ssf1-Rrp15 heterodimer, which stabilizes the junction between domains I and VI in the later State 2 of the nucleolar pre-60S^17^.

Thus, by various modes of action, including steric hindrance and pre-rRNA remodeling, Noc1-Noc2 serves as a central nexus that stabilizes a co-transcriptional pre-rRNA folding intermediate while delaying subsequent post-transcriptional maturation events until transcription of the entire large subunit pre-rRNA is completed.

Previous biochemical data has shown that the transcription of the LSU pre-rRNA precedes co-transcriptional cleavage at site A2, suggesting a ribosome assembly checkpoint ^2^. Consistent with this data, the structure of the Noc1-Noc2 RNP indicates that the assembly of this particle likely serves as checkpoint since the Noc1-Noc2 dimer interrogates the formation of root helices that form the core of the nascent LSU, encapsulates helix 2, and stabilizes the 5’ end of the 5.8S rRNA (**Figs. 1, 2a**). To probe this ribosome assembly checkpoint, we designed assays to interrogate pre-rRNA processing *in vivo* and protein composition of different pre-rRNP species (**Fig. 3**). To investigate the ribosome assembly checkpoint *in vivo*, we recapitulated co-transcriptional ribosome assembly by transforming yeast strains with plasmids containing unique rDNAs that differ in their ability to form helix 2 (**Fig. 3a**). Since helix 2 is bound directly by Noc1-Noc2, we hypothesized that an inability to form a base-paired helix 2 should affect both pre-rRNA processing and the association of early assembly factors such as the Noc1-Noc2 complex. To reflect endogenous co-transcriptional ribosome assembly, the designed plasmids encoded for an entire small subunit precursor (5’ ETS and 18S rRNA) followed by ITS1 and variants of LSU RNA elements that can be distinguished from the endogenous ribosomal subunits by virtue of distinct probes for Northern blotting (**Fig. 3a**). The structural integrity of helix 2 is key to this ribosome assembly checkpoint and pre-rRNAs that included both strands of helix 2 (domain II, domain I, and domain I-424nt) were detected as unprocessed transcripts (**Fig. 3b**) and could be further processed at site A2 to give rise to stable LSU precursors (**Fig. 3c, d**). By contrast, pre-rRNAs where helix 2 was disrupted either by truncation to remove one strand of helix 2 (domain I-404nt) or mutation of either strand of helix 2 (domain II-5.8S mutant, domain II-25S mutant) were less stable (**Fig. 3c, d**). Strikingly, compensatory mutations that restore the RNA duplex within helix 2 (domain II-rescue mutant, **Fig. 3a**) restore a stable precursor (**Fig. 3b**) that is effectively processed at site A2 (**Fig. 3c-e)**.

**Figure 2.**
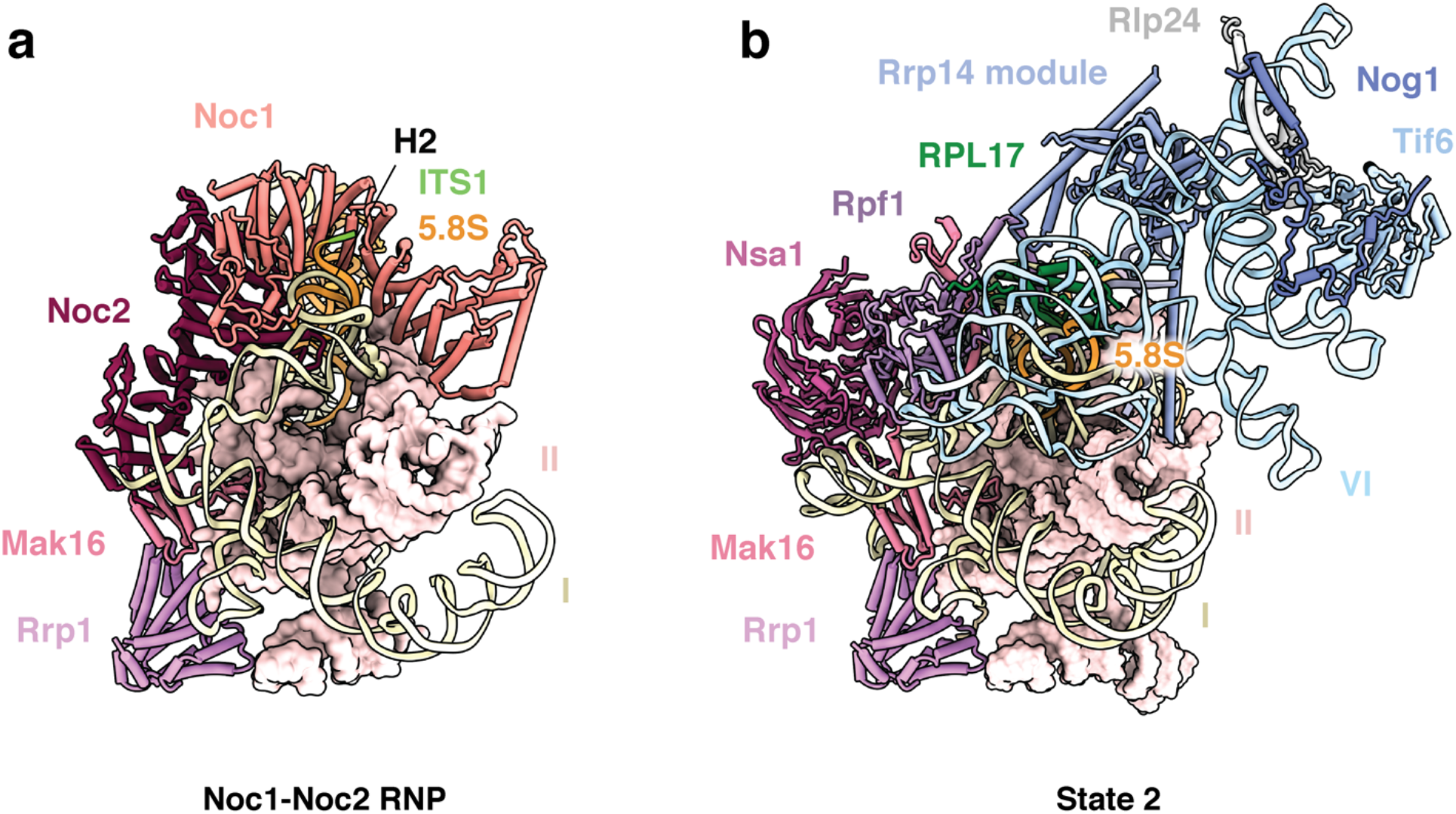
The Noc1-Noc2 complex controls early large subunit assembly. (**a**) Cartoon representation of color-coded assembly factors chaperoning domains I and II of the Noc1-Noc2 RNP. Domain II is shown in surface representation and root helix 2 (H2) and ITS1 are indicated. (**b**) A comparative view of State 2 (PDB: 6C0F, superimposed on Rrp1) with cartoon representation of color-coded assembly factors chaperoning domains I,II and VI. Domain II is shown in surface representation and ribosomal protein RPL17 is indicated in green.

**Figure 3.**
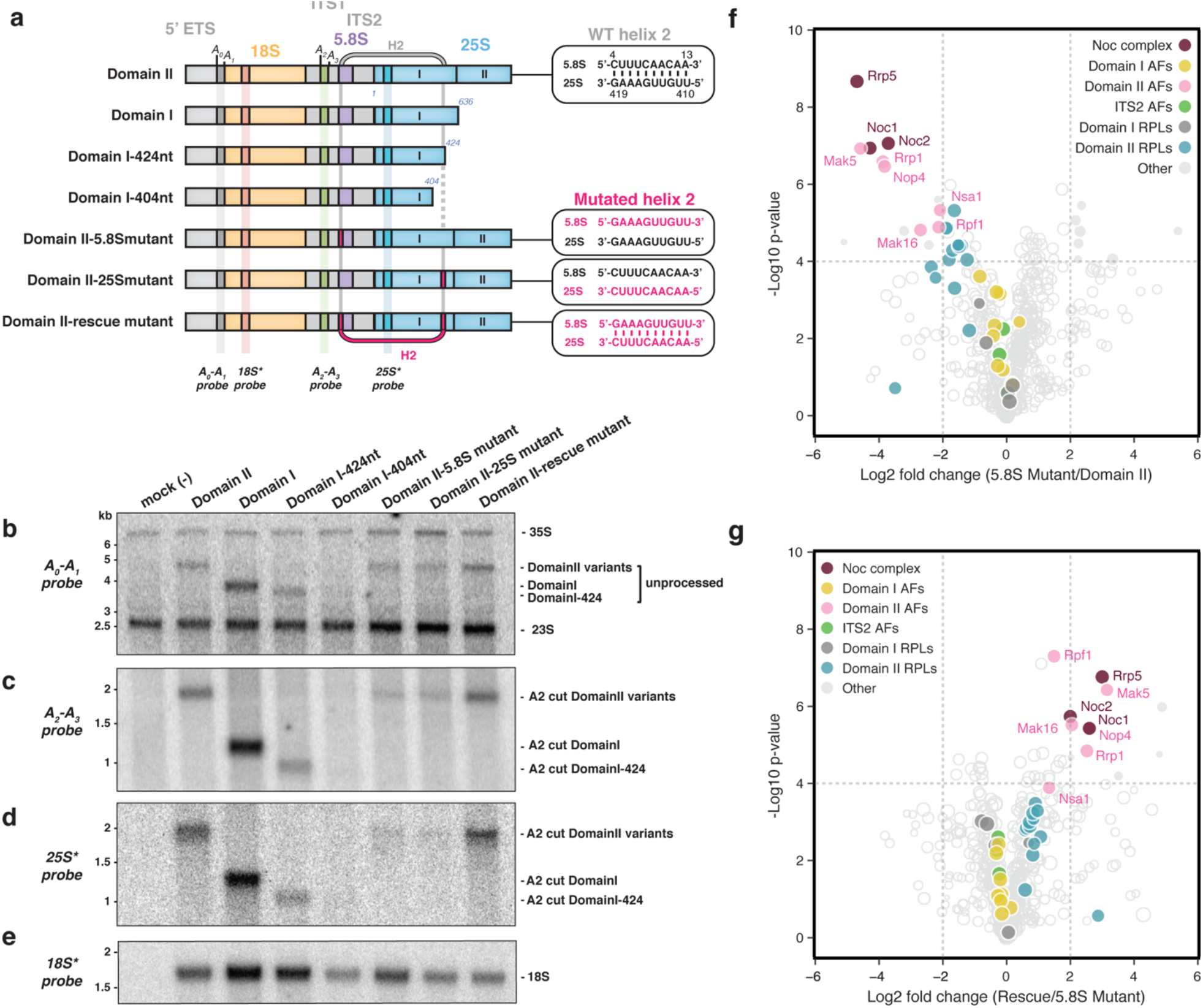
Binding of Noc1-Noc2 verifies correct formation of helix 2. **(a)** A schematic of rRNA mimics (Domain II, Domain I, helix 2 truncations and mutants) expressed in yeast and analyzed by northern blot. WT and mutated helix 2 (H2) are depicted in gray and magenta respectively with a dotted line representing missing fragment of H2. **(b, c, d, e)** Northern blot analysis of total RNA extracted from whole cells transformed with the large subunit rRNA variants shown in (a), (b) probed with A0-A1 probe, (c) A2-A3 probe, (d) 25S* (specifically inserted) probe, and (e) 18S* (specifically inserted) probe. **(f, g)** Comparative mass spectrometry analysis of proteins associated with large subunit rRNA variants starting at site A2 and terminating with Domain II. (f) Comparison of proteins associated with Domain II-5.8S mutant and Domain II. (g) Comparison of proteins associated with Domain II-rescue mutant and Domain II-5.8S mutant. Log2 fold changes in protein abundance are expressed as mean LFQ values of 3 replicates. P-values were determined by Student’s t test and P-value of 0.0001 was chosen as cut-off.

To investigate whether the structural integrity of the RNA duplex of helix 2 is important for cooperative stabilization by proximal assembly factors, we performed comparative mass spectrometry with designed LSU rRNA precursors (**Fig. 3f, g**). Here, a single strand mutant of helix 2 in the 5.8S rRNA (domain II-5.8S mutant) selectively reduced the levels of proteins associated with domain II. The most affected proteins were proteins directly associated with helix 2 (Noc1, Noc2 and Rrp5) followed by proximal assembly factors associated with Mak16 (Rpf1, Rrp1 and Nsa1) and domain II associated ribosomal proteins, while ribosomal proteins associated with domain I were unaffected (**Fig. 3f, Extended Data Fig. 10**). By contrast, compensatory mutations within helix 2 that restore the RNA duplex (domain II-rescue mutant) selectively re-established binding of direct interactors of helix 2 (Noc1, Noc2, Rrp5), as well as proximal assembly factors associated with Mak16 (Rpf1, Rrp1 and Nsa1) and domain II associated ribosomal proteins (**Fig. 3g, Extended Data Fig. 10**).

These data strongly support a model in which the formation of the nascent Noc1-Noc2 RNP constitutes an assembly checkpoint where local stabilization of core root helices (helix 2, 25 and 26) is communicated through the cooperative association of both assembly factors and ribosomal proteins. An inability to form a key root helix within the pre-rRNA (such as helix 2) prevents co-operative stabilization of the precursor by ribosomal proteins and assembly factors (such as Noc1-Noc2), thereby inhibiting downstream processing steps.

The combined structural and biochemical data presented here suggest that the coupled assembly of eukaryotic small and large ribosomal subunits can be described by the following model (**Fig. 4**): After the co-transcriptional assembly of the small subunit processome (**Fig. 4a**), the transcription of ITS1, the 5.8S rRNA, ITS2 and most of the 25S rRNA domain I enables the formation of helix 2, which is sufficient to stabilize a co-transcriptional LSU assembly intermediate that can undergo cleavage at site A2 (**Fig. 4b**). The subsequent transcription of domain II generates the co-transcriptional assembly intermediate revealed in this study, that also contains the tethered assembly factor complex Mak16-Rpf1-Nsa1 and remains a part of the nascent pre-rRNP visualized in Miller spreads until rDNA transcription is completed (**Fig. 4a, c**). Following the complete transcription of the LSU pre-rRNA, post-transcriptional assembly events occur, where the departure of the Noc1-Noc2 complex, the integration of domain VI and the stable incorporation of Rpf1-Nsa1 lead to the formation of State 2 of the nucleolar pre-60S in which domains I and II are joined (**Fig. 4d**). This is then followed by the association of the Noc2-Noc3 complex, which probes the polypeptide exit tunnel in State E (**Fig. 4e**). These mechanisms form the basis for the highly controlled initial stages of both co- and early post-transcriptional assembly and processing of the eukaryotic LSU.

**Figure 4.**
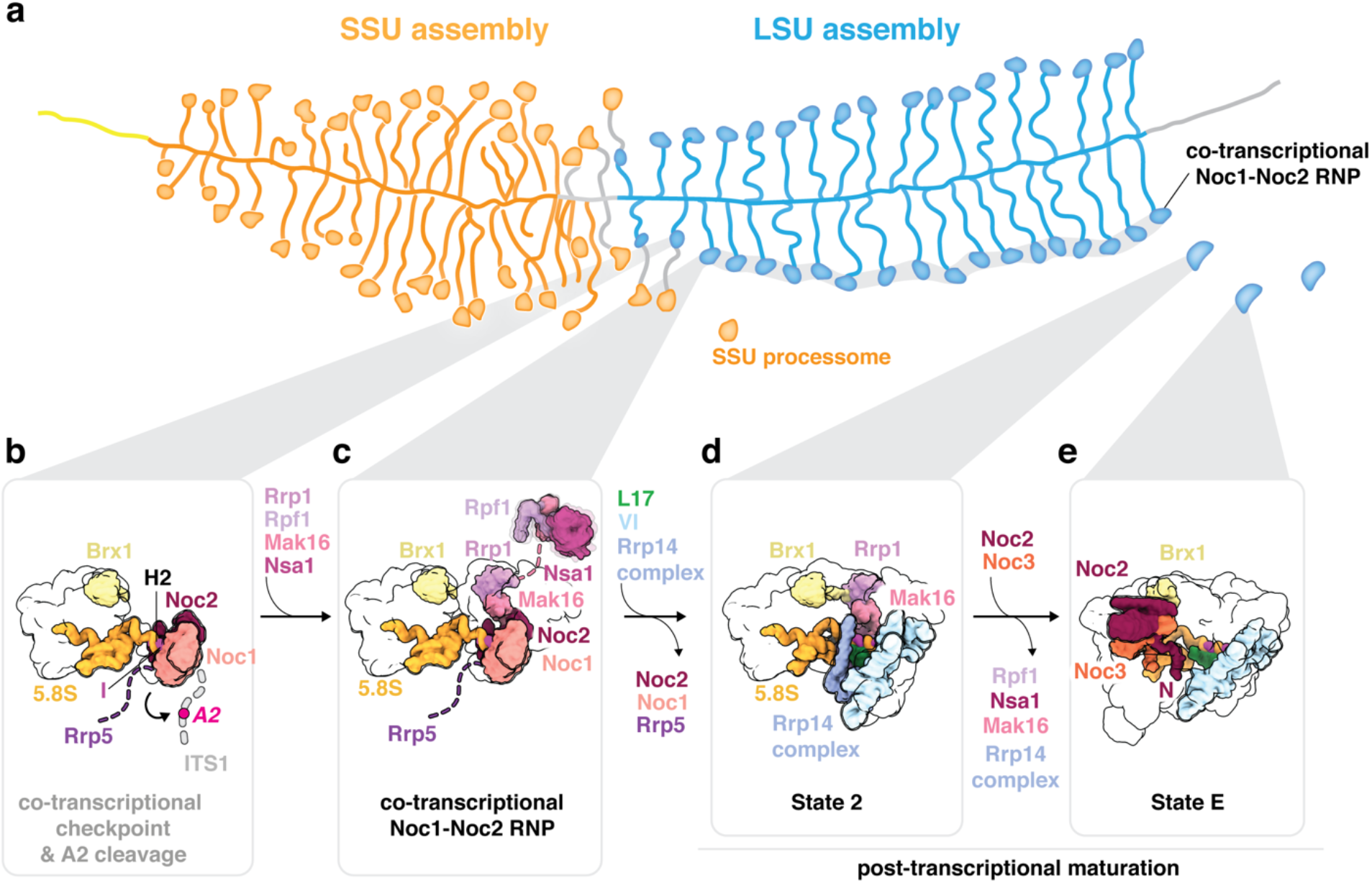
A model for quality control during nucleolar LSU assembly. **(a)** Representation of a Miller spread where transcription of rDNA results in nascent pre-rRNA decorated by protein factors that chaperone the pre-rRNA into folded structures or terminal knobs. **(b)** Schematic model for how upon transcription of domain I of the LSU, co-transcriptional assembly occurs with Noc1-Noc2-Rrp5 chaperoning the rRNA around helix 2, mediating a critical checkpoint and regulating the cleavage of the rRNA at site A2. **(c)** Upon passing of this quality check, the Noc1-Noc2 RNP is formed with domains I and II of the 25S rRNA incorporated (this study, PDB 8E5T). **(d)** Post-transcriptional maturation continues with State 2 as more ribosomal proteins join and further compaction of rRNA domains occur^17^. **(e)** Noc2-Noc3 RNP (State E) represent the later stages of nucleolar maturation with most of the 25S rRNA domains ordered (this study, EMD-27910).

Our structural and functional data shed light on the earliest events during co-transcriptional large ribosomal subunit formation. Together, these data provide a framework to explain a co-transcriptional ribosome assembly checkpoint centered around the 5’ end of nascent LSU precursors in the context of Miller spreads.

