## Supplementary Information for "A co-transcriptional ribosome assembly checkpoint controls nascent large ribosomal subunit maturation"

### METHODS

Cloning of the MS2-tagged 25S rRNA and the MS2-3c-GFP construct

The large subunit rDNA, beginning at the A2 site in ITS1 and terminating after Domain VI of the 25S, was cloned from the rDNA locus of the *Saccharomyces cerevisiae* strain BY4741 into a derivative of the pESC_URA vector (Agilent Technologies). The resulting rRNA mimic (A2-DomainVI) was tagged with five MS2-aptamer stem-loops at its 3' end and cloned downstream of a gal1 promoter and upstream of a CYC terminator. An adapted MS2-coat protein fused to an N-terminal nuclear localization signal (NLS), a hemagglutinin (HA) tag and a C-terminal 3C-protease-cleavable GFP (NLS-HA-MS2-3C-GFP) was cloned into a modified pESC plasmid suitable for genome integration in yeast, under a gal10 promoter with G418 resistance.

Expression and purification of pre-60S particles

The pESC plasmid containing the MS2-3C-GFP under the gal10 promoter was transformed into the *S. cerevisiae* BY4741 strain for integration by selecting for G418 resistance. The resulting yeast strain with MS2-3C-GFP integrated, was then subjected to endogenous tagging of a selected assembly factor – Noc1 (Mak21), Noc2, or Cic1 with a C-terminal streptavidin-binding-peptide (sbp) tag using ClonNAT (Nourseothricin) selection. The subsequent strain containing both the integrated galactose-inducible MS2-3C-GFP and the C-terminal sbp tagged assembly factor was transformed with the pESC_URA plasmid containing the pre-60S rDNA for transient expression.

Yeast cultures were grown in Ura- synthetic drop-out (SD) media containing 2% galactose (w/v) at 30 ºC for 16-18 hours, reaching saturation (OD 5-6). Cells were then harvested by centrifugation at 3000 x *g* for 10 minutes at 4 ºC, resulting in a cell mass of 20-30 grams. The cell pellet was washed with ice cold $ddH_2O$ twice, followed by a wash with $ddH_2O$ containing protease inhibitors (E64, Pepstatin, PMSF). Washed cells were immediately flash frozen in liquid nitrogen and lysed by 4 cycles of cryogenic grinding using a Retsch Planetary Ball Mill PM100. The freshly ground yeast powder was resuspended by vortexing in buffer A (50 mM Tris-HCl, pH 7.6 (20 °C), 150 mM NaCl, 5mM $MgCl_2$, 1 mM DTT, 0.1% Triton-X100, PMSF, Pepstatin, E-64), followed by centrifugation at 4 °C, 40,000 x *g* for 30 min to remove the insoluble component. The supernatant was then incubated with anti-GFP nanobody beads (Chromotek) for 3 hours at 4 °C, with gentle agitation. The beads were washed three times in ice-cold buffer A and once in buffer B (50 mM Tris-HCl pH 7.6 (20 °C), 150 mM NaCl, 5 mM $MgCl_2$, 1 mM DTT), before the bound proteins were eluted with 3C-protease cleavage for 1 hour at 4 °C. The eluate in buffer B was then applied to NHS-sepharose beads (Sigma) coupled with streptavidin for 1 hour at 4 °C with agitation. The pre-60S bound streptavidin beads were washed once with buffer B, before release from beads using buffer B containing 5mM d-Biotin. The eluted sample typically measured an absorbance at 260 nm ($A_{260}$) of 1.0 to 2.5 mAU (Nanodrop 2000, Thermo Scientific) (Extended Data Fig. 1). The quality of the sample was initially judged by SDS-PAGE and negative stain electron microscopy (EM) and then used for preparing cryo-EM grids. Mass-spectrometry analysis of the sample allowed for identification of expected protein components in the purified complex.

Cryo-EM sample and grid preparation

Cryo-EM grids were prepared from multiple purifications for the 8 data sets obtained

(DS1-DS8) for the Noc1 particle. The eluate in buffer B (above) was supplemented with 0.1% Triton X-100 (final concentration) prior to grid preparation. Copper grids of 400 mesh with lacey carbon and an ultra-thin carbon support film were used (Ted Pella Inc, product no. 01824) for data collection. A volume of 3 to 4 µl of sample ($A_{260}$ of ~1.5) was applied onto glow-discharged grids, incubated for 30 seconds and plunged into liquid ethane using a Vitrobot Mark IV robot (FEI Company) (95 % humidity, blot force of 2-4 and blot time 3.5-4 s). Cryo-EM grids for the Noc2-Noc3 particle were prepared in the same manner.

### Cryo-EM data collection and image processing

**Noc1-Noc2 RNP:** A total of 18,046 micrographs were obtained over eight data collections on a Titan Krios (Thermo Fisher), at 300 kV with a K2 Summit detector (Gatan, Inc.). SerialEM [24] was employed for data acquisition using a defocus range of 1.0- 3.0 µm with a pixel size of 1.3 Å. Super-resolution movies (pixel size 0.65 Å), with 32 frames were collected using a total dose of 8 electrons per pixel per second with an exposure time of 8 seconds and a total dose of 37.9 electrons per Å$^2$ (Supplementary Table 1).

Upon data collection, the movies were gain corrected, dose weighted, aligned, and binned to pixel size of 1.3 Å using RELION 3.0's implementation of a Motioncor2-like algorithm, and the contrast transfer function (CTF) was estimated using CTFFIND 4.1 [25] within RELION [26]. Corrected and aligned micrographs were subjected to automated particle picking by CrYOLO [27], which resulted in a total of 1,878,650 particles from all 8 data sets combined. Particle coordinates were then imported into RELION for subsequent steps. Particles were extracted with a box size of 220 pixels (pixel size of 2.6 Å/px), and 2D-classified separately for each individual data set. After 2D classification, bad classes were removed and selected particles from each data set were combined resulting in a total particle stack of 977,232 particles. The particles were subjected to a round of Bayesian polishing in RELION, followed by global search 3D classification into five classes (K=5) using an initial 3D model obtained from an *ab-initio* reconstruction from DS1 in RELION, low-pass filtered to 40 Å. The best two classes from this 3D classification were selected and their particles were re-extracted with a box size 440 pixels (pixel size of 1.3 Å/px). A combined total of 556,013 particles were used for a consensus 3D refinement with a solvent mask containing the entire particle, resulting in an overall resolution of 6.9 Å.

Due to observed inherent flexibility of the particle between domains I and II of the 25SrRNA, the particles were subjected to particle subtraction in RELION to remove signal from each domain respectively, and process them separately, in order to improve the quality and resolution of the density. Upon subtraction of the signal using individual masks around each domain, a 3D refinement followed by a round of 3D classification without image alignment was performed, first for domain II (K=8). The best class was selected, and checked for duplicates, resulting in a final particle stack of 158,915 particles. The particles were then subjected to 3 rounds of CTF refinement and Bayesian polishing, resulting in a final reconstruction of domain II at an overall resolution of 3.6 Å. This particle stack was then reverted back to its original form and the signal from domain II was removed providing a starting point to obtain the domain I reconstruction. The particles were 3D classified without image alignment, (K=8), and three of the most similar classes were selected and combined, resulting in a total particle number of 49,406. These particles were then further classified with alignment (K=5), and a single class was chosen with 17,104 particles. After 3 rounds of CTF refinement and particle polishing in RELION, a final reconstruction of domain I at a resolution of 4.72 Å was obtained (Extended Data Fig. 2). The local resolution of the maps was calculated with blocres [28] within cryoSPARC [29] (Extended Data Fig. 3). All reported resolutions are based on the gold standard FSC-0.143 criterion and FSC-curves were corrected using high-resolution noise substitution methods in RELION 3.0. Composite map for the entire complex was generated using phenix.combine_focused_maps [30]. The same program was used to generate composite half maps that were used to calculate the overall resolution of the reconstruction at 4.0 Å.

**Noc2-Noc3 RNP:** A total of 8,249 micrographs from 2 data collections were obtained, on the Titan Krios (Thermo) using the same grid preparation and collection parameters as was described above for the Noc1-particle. Particles were picked using CrYOLO, resulting in 253,724 and 416,087 particles from each data set respectively. The extracted particles (box size 440 pixels) were subjected to a round of 2D classification and 3D classification with alignment (K=5) separately, to obtain one good 3D class from each data set which were then combined to obtain a final particle stack of 75,358 particles. The particles were then 3D-refined and put through two rounds of CTF refinement and Bayesian polishing resulting in a final reconstruction of the Noc2-Noc3 particle at 3.69 Å resolution. The local resolution of the maps was calculated with blocres within cryoSPARC (Extended Data Fig. 3). All reported resolutions are based on the gold standard FSC-0.143 criterion and FSC-curves were corrected using high-resolution noise substitution methods in RELION 3.0.

Model building and refinement

Using the structure of the *S. cerevisiae* early nucleolar pre-60S particle (PDB 6C0F) [17] as reference, common assembly factors and ribosomal proteins were manually located and fitted into the density. New assembly factors Noc1 and Noc2 were initially built *de novo* while using models predicted by AlphaFold [31] subsequently as references, and remodeled segments of the 25S and 5.8S rRNA were also built *de novo* into the domain II density. Rigid body docking was used for domain I factors and rRNA. Model building was performed with COOT [32]. An annotated list of individual protein IDs, reference models and corresponding maps used for building, can be found in Supplementary Table 2. The final model was refined against the composite map in PHENIX with phenix.real_space_refine using secondary structure restraints for proteins and RNAs [30]. Refinement and model statistics can be found in Supplementary Table 1. All map and model analyses and illustrations were made using USCF ChimeraX (version 1.2.5) [33] and PyMOL Molecular Graphics System (Version 2.3.5 Schrödinger, LLC).

RNA extraction and Northern blotting

The rRNA mimics were purified as described above and RNA was extracted from the final eluate with 1 mL TRIzol (Life Technologies) according to the manufacturer’s instructions. The whole cell RNA was extracted by resuspending 0.2 g of yeast cells in 200 µL of TRIzol followed by lysis by bead beating for the total time of 10 mins. The total volume of TRIzol was then brought to 1 mL and the extraction continued according to the manufacturer’s instructions. 1.0 µg of isolated rRNA was separated on a denaturing 1.2% Formaldehyde-Agarose gel (SeaKem LE, Lonza). After staining the gel in 1X SYBR Green II (Lonza)

$ddH_2O$ solution (pH 7.5) for 30 min, RNA species were visualized with a Gel Doc EZ Imager (Bio-Rad) (Extended Data Fig. 1) and then transferred onto a cationized nylon membrane (Zeta-Probe GT, Bio-Rad) using downward capillary transfer. The RNA was cross-linked to the membrane for Northern blot analysis by UV irradiation at 254 nm with a total exposure of 120 milli- joules/$cm^2$ in a UV Stratalinker 2400 (Stratagene). Cross-linked membranes were initially stained with methylene blue dye to visualize quality of transfer and then were incubated with hybridization buffer (750 mM NaCl, 75 mM trisodium citrate, 1% (w/v) SDS, 10% (w/v) dextran sulfate, 25% (v/v) formamide) at 65°C for 30 min prior to addition of γ-$^{32}$P-end-labeled DNA oligo nucleotide probe.

Used oligonucleotide probe sequences:

25S probe*: AGGTACACTCGAGAGCTTCA

18S probe*: CGAGGATCGAGGCTTT

5'ETS ($A_0$-$A_1$) probe: CCCACCTATTCCCTCTTGC

ITS1 ($A_2$-$A_3$) probe: TGTTACCTCTGGGCCCCGATTG

The probe was hybridized for 1 hour at 65 °C and then overnight at 37 °C. Membranes were washed once with wash buffer 1 (300 mM NaCl, 30 mM trisodium citrate, 1% (w/v) SDS) and once with wash buffer 2 (30 mM NaCl, 3 mM trisodium citrate, 1% (w/v) SDS) for 20 min each at 45°C. Radioactive signal was detected by exposure of the washed membranes to a storage phosphor screen which was scanned with a Typhoon 9400 variable-mode imager (GE Healthcare).

Mass spectrometry and Comparative data analysis

Purified RNP samples were dried and dissolved in 8 M urea/0.1 M ammonium bicarbonate/10 mM DTT. After reduction, cysteines were alkylated in 30 mM iodoacetamide (Sigma). Proteins were digested with LysC (Endoproteinase LysC, Wako Chemicals) in less than 4 M urea followed by trypsination (Trypsin Gold, Promega) in less than 2 M urea. Digestions were halted by adding TFA and digests were desalted [34] and analyzed by reversed phase nano-LC-MS/MS using a Fusion Lumos (Thermo Scientific). Data were quantified and searched against the *S. cerevisiae* Uniprot protein database (2019) concatenated with the MS2-protein sequence and common contaminations. For the search and quantitation, MaxQuant v. 2.0.3.0 [35] was used. Oxidation of methionine and protein N-terminal acetylation were allowed as variable modifications and all cysteines were treated as being carbamidomethylated. The ‘match between runs’ option was enabled, and false discovery rates for proteins and peptides were set to 1% and 2% respectively.

Protein abundances were expressed as LFQ (label free quantification) values. Data were analyzed using Perseus (v.1.6.10.50) [36]. In short, LFQ values were LOG2 transformed followed by a filtering requiring that a protein must be matched in all 3 replicates for at least one condition. A student t-test was carried out to confirm statistical significance of data.

**Data and materials availability**: The cryo-EM maps and atomic models have been deposited in the Electron Microscopy Data Bank (EMDB) and the Protein Data Bank (PDB): Noc1-Noc2 RNP (EMDB-27919, PDB 8E5T) and Noc2-Noc3 RNP (EMDB-27910). Raw unaligned multi-frame movies and aligned micrographs of the Noc1-Noc2 RNP have been deposited in the Electron Microscopy Public Image Archive (EMPIAR-XXXX). Materials are available from S.K upon reasonable request under a material transfer agreement with the Rockefeller University.

**Acknowledgments**

We would like to thank M. Ebrahim, J. Sotiris, and H. Ng at the Evelyn Gruss Lipper Cryo-Electron Microscopy Resource Center at The Rockefeller University for assistance with grid screening and data collection. Mass spectrometry data was generated by the Proteomics Resource Center at The Rockefeller University (RRID:SCR_017797) using instrumentation funded by the Sohn Conferences Foundation and the Loena M. and Harry B. Helmsley Charitable Trust, and we particularly thank H. Molina, S. Heissel, and C. Peralta for assistance. We thank F. Stengel and C. Sailer for sharing cross-linking and mass spectrometry data ahead of publication, O. Buzovetsky for assistance with model building, and members of the Klinge laboratory for critical reading of the manuscript.

**Author information:**

Laboratory of Protein and Nucleic Acid Chemistry, The Rockefeller University, New York, NY, USA

Zahra A. Sanghai, Rafal Piwowarczyk, Arnaud Vanden Broeck and Sebastian Klinge

**Funding**: This work was supported by funding from the G. Harold and Leila Y. Mathers Foundation (MF-2104-01554) and the National Institutes of Health grant GM143181 to S.K.

**Author contributions**: Z.A.S and S.K conceived the study, Z.A.S performed RNA analysis, cryo-EM sample preparation, data collection, processing, and model building. R.P. performed biochemical experiments and mass spectrometry analysis, A.V.B. performed cryo-EM processing and model building, and all authors wrote and edited the manuscript.

**Competing interests**: none declared.

**Supplementary Information is available for this paper.**

### Extended Data

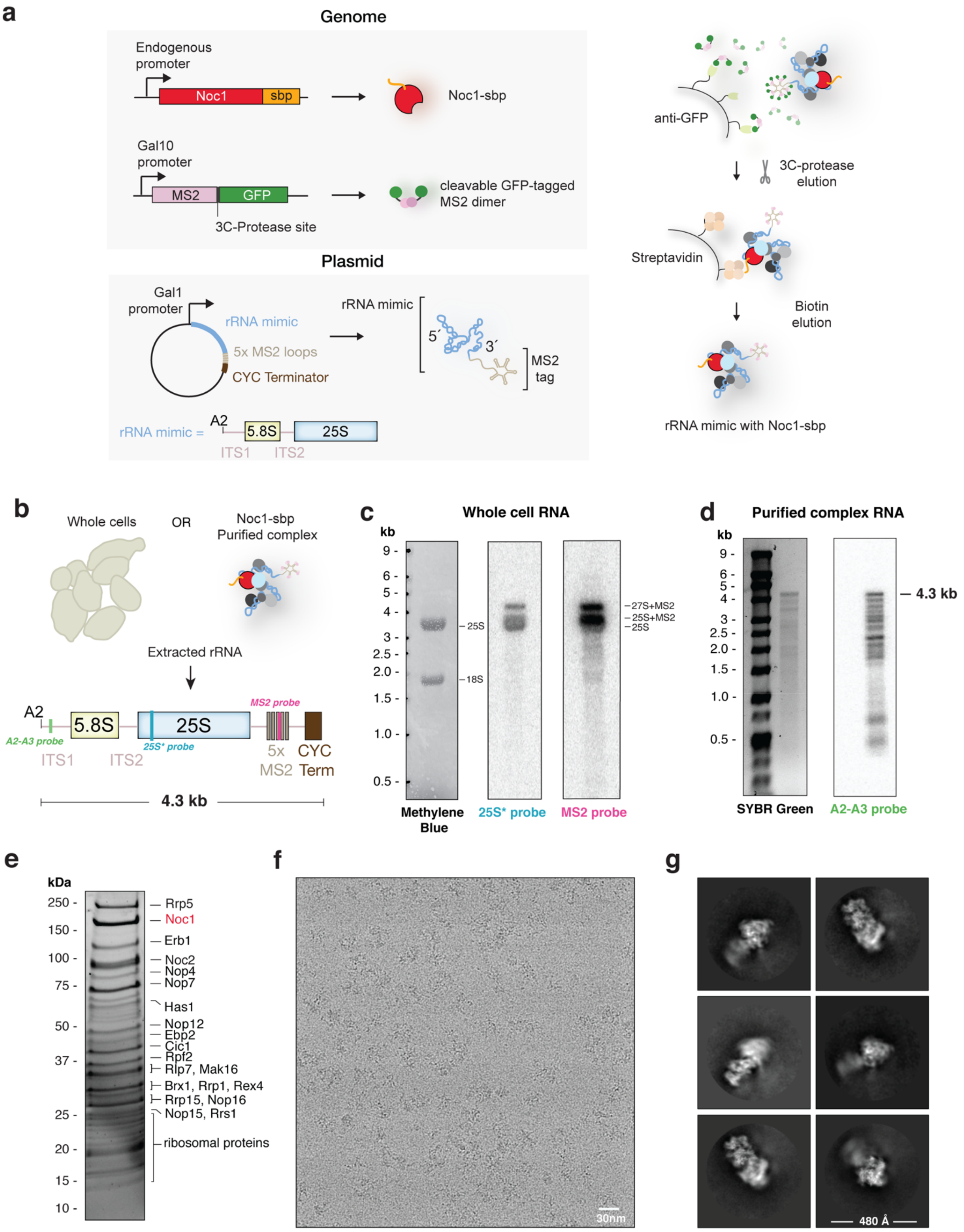

**Extended Data Fig. 1: Noc1-Noc2 RNP expression and purification scheme and cryo-EM images.** (**a**) Genomic integration and composition of the plasmid containing the rRNA mimic used for Noc1-Noc2 RNP expression in yeast, followed by schematic of the two-step purification of the Noc1-Noc2 RNP (**b**) Schematic of RNA extraction from whole cells (see panel c) and the purified Noc1-Noc2 RNP (see panel d). (**c**) Methylene blue-stained RNA gel and northern blot of RNA extracted from whole cells containing the Noc1-Noc2 RNP. (**d**) SYBR Green-stained RNA gel and northern blot of RNA extracted from purified Noc1-Noc2 RNP. **(e)** Representative SYPRO Ruby stained SDS-PAGE of purified fraction containing the Noc1-Noc2 RNP pre-ribosomal particles. Molecular weight markers are indicated on the left (MW) and bait protein (Noc1; red) and other bands were identified by LC-MS/MS analysis of eluate. (**f**) Representative motion-corrected cryo-EM micrograph of the Noc1-Noc2 RNP particle sample. (**g**) Six most-populated 2D class averages (1.3 Å/px), extracted box size 440 pixels, mask diameter 480 Å.

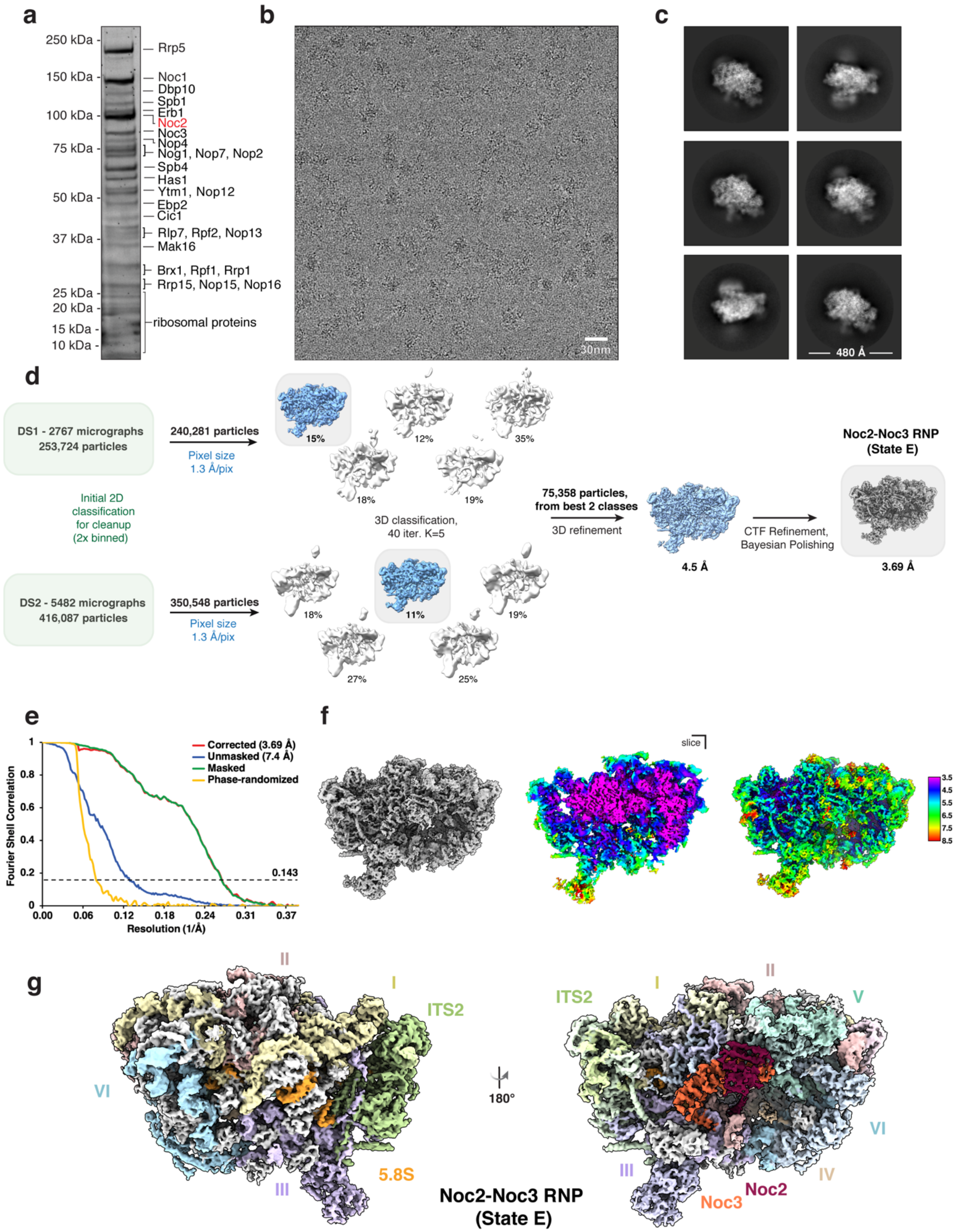
a
250 kDa
150 kDa
100 kDa
75 kDa
50 kDa
37 kDa
25 kDa
20 kDa
15 kDa
10 kDa
Rrp5
Noc1
Dbp10
Spb1
Erb1
Noc2
Noc3
Nop4
Nog1, Nop7, Nop2
Spb4
Has1
Ytm1, Nop12
Ebp2
Cic1
Rlp7, Rpf2, Nop13
Mak16
Brx1, Rpf1, Rrp1
Rrp15, Nop15, Nop16
ribosomal proteins
b
30nm
c
480 Å
d
DS1 - 2767 micrographs
253,724 particles
240,281 particles
Pixel size
1.3 Å/pix
Initial 2D
classification
for cleanup
(2x binned)
DS2 - 5482 micrographs
416,087 particles
350,548 particles
Pixel size
1.3 Å/pix
15%
12%
35%
18%
19%
3D classification,
40 iter. K=5
18%
11%
19%
27%
25%
75,358 particles,
from best 2 classes
3D refinement
4.5 Å
CTF Refinement,
Bayesian Polishing
Noc2-Noc3 RNP
(State E)
3.69 Å
e
Corrected (3.69 Å)
Unmasked (7.4 Å)
Masked
Phase-randomized
Fourier Shell Correlation
0.143
Resolution (1/Å)
f
slice
g
II
I
ITS2
VI
III
5.8S
180°
V
IV
Noc3
Noc2
Noc2-Noc3 RNP
(State E)

**Extended Data Fig. 2. Purification and structure determination of the Noc2-Noc3 RNP (State E).**

**(a)** Representative SYPRO Ruby stained SDS-PAGE of purified fraction containing the Noc2-Noc3 RNP (State E) pre-ribosomal particles. Molecular weight markers are indicated on the left (MW) and bait protein (Noc2; red) and other bands were identified by LC-MS/MS analysis of the eluate. (**b**) Representative motion-corrected cryo-EM micrograph of the Noc2-Noc3 RNP particle sample. (**c**) Six most populated 2D class averages (1.3 Å/px), box size 440 pixels, mask diameter 480 Å. **(d)** Cryo-EM data acquisition and processing workflow to obtain the final reconstruction of the Noc2-Noc3 RNP at a resolution of 3.69 Å. **(e)** Solvent-corrected FSC curves for the Noc2-Noc3 RNP, calculated in Relion 3.0, with reported resolutions determined at FSC=0.143 upon post-processing. **(f)** Local resolution estimation displayed in rainbow color on the cryo-EM map of the Noc2-Noc3 RNP. **(g)** Final cryo-EM reconstruction of the Noc2-Noc3 RNP with rRNA domains, assembly factors and ribosomal proteins (in gray) identified and colored.

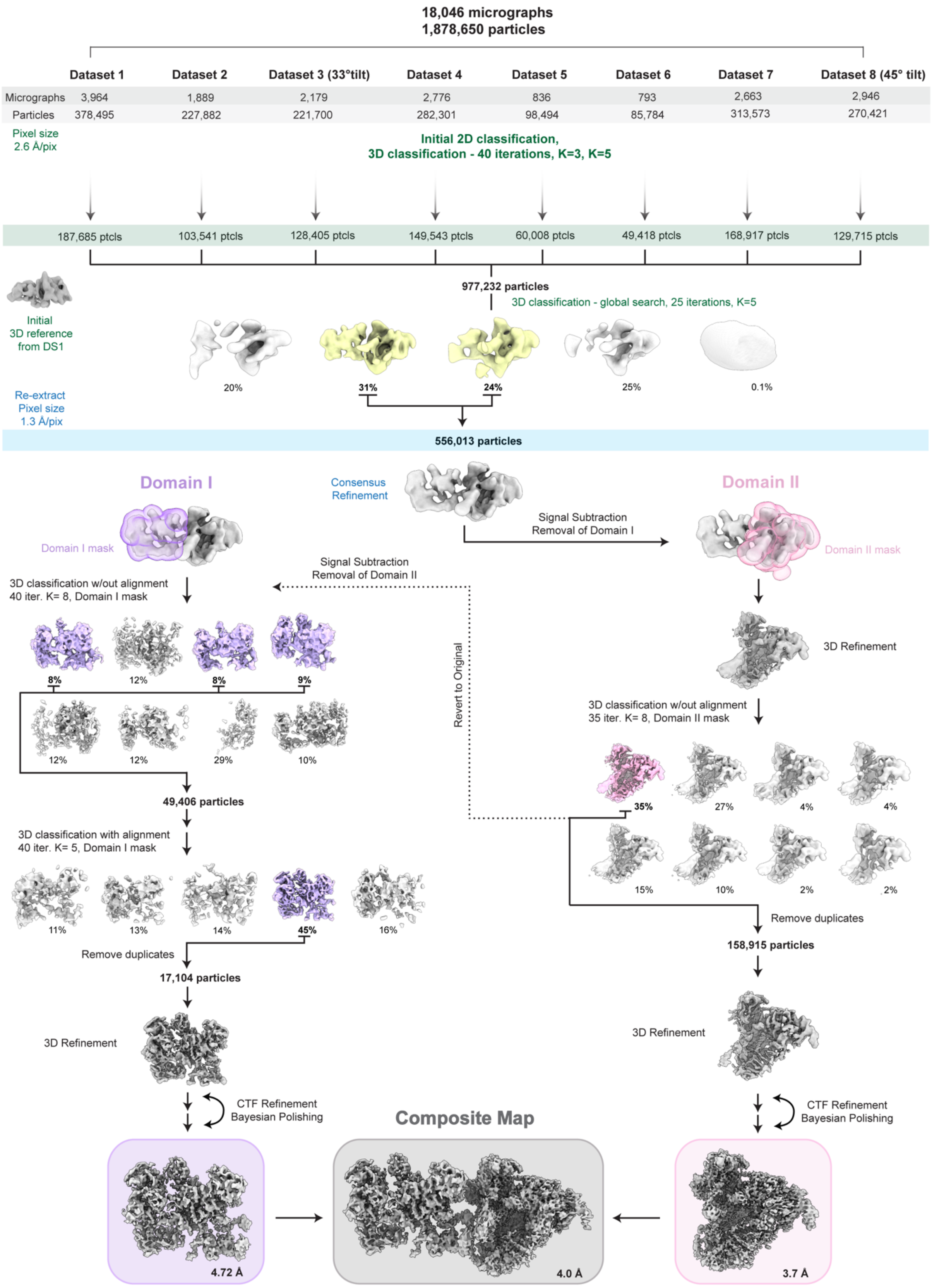
18,046 micrographs
1,878,650 particles
Dataset 1
Dataset 2
Dataset 3 (33°tilt)
Dataset 4
Dataset 5
Dataset 6
Dataset 7
Dataset 8 (45° tilt)
Micrographs 3,964 1,889 2,179 2,776 836 793 2,663 2,946
Particles 378,495 227,882 221,700 282,301 98,494 85,784 313,573 270,421
Pixel size 2.6 Å/pix
Initial 2D classification, 3D classification - 40 iterations, K=3, K=5
187,685 ptcls
103,541 ptcls
128,405 ptcls
149,543 ptcls
60,008 ptcls
49,418 ptcls
168,917 ptcls
129,715 ptcls
977,232 particles
3D classification - global search, 25 iterations, K=5
Initial 3D reference from DS1
20%
31%
24%
25%
0.1%
Re-extract Pixel size 1.3 Å/pix
556,013 particles
Domain I
Consensus Refinement
Domain II
Signal Subtraction Removal of Domain I
Domain I mask
Domain II mask
Signal Subtraction Removal of Domain II
3D classification w/out alignment 40 iter. K= 8, Domain I mask
8%
12%
8%
9%
12%
12%
29%
10%
Revert to Original
3D Refinement
3D classification w/out alignment 35 iter. K= 8, Domain II mask
35%
27%
4%
4%
15%
10%
2%
2%
49,406 particles
3D classification with alignment 40 iter. K= 5, Domain I mask
11%
13%
14%
45%
16%
Remove duplicates
17,104 particles
Remove duplicates
158,915 particles
3D Refinement
3D Refinement
CTF Refinement Bayesian Polishing
Composite Map
CTF Refinement Bayesian Polishing
4.72 Å
4.0 Å
3.7 Å

**Extended Data Fig. 3. Data collection and structure determination of the Noc1-Noc2 RNP.**

Cryo-EM data acquisition and processing workflow for the eight data sets collected to obtain the final reconstruction of the Noc1-Noc2 RNP.

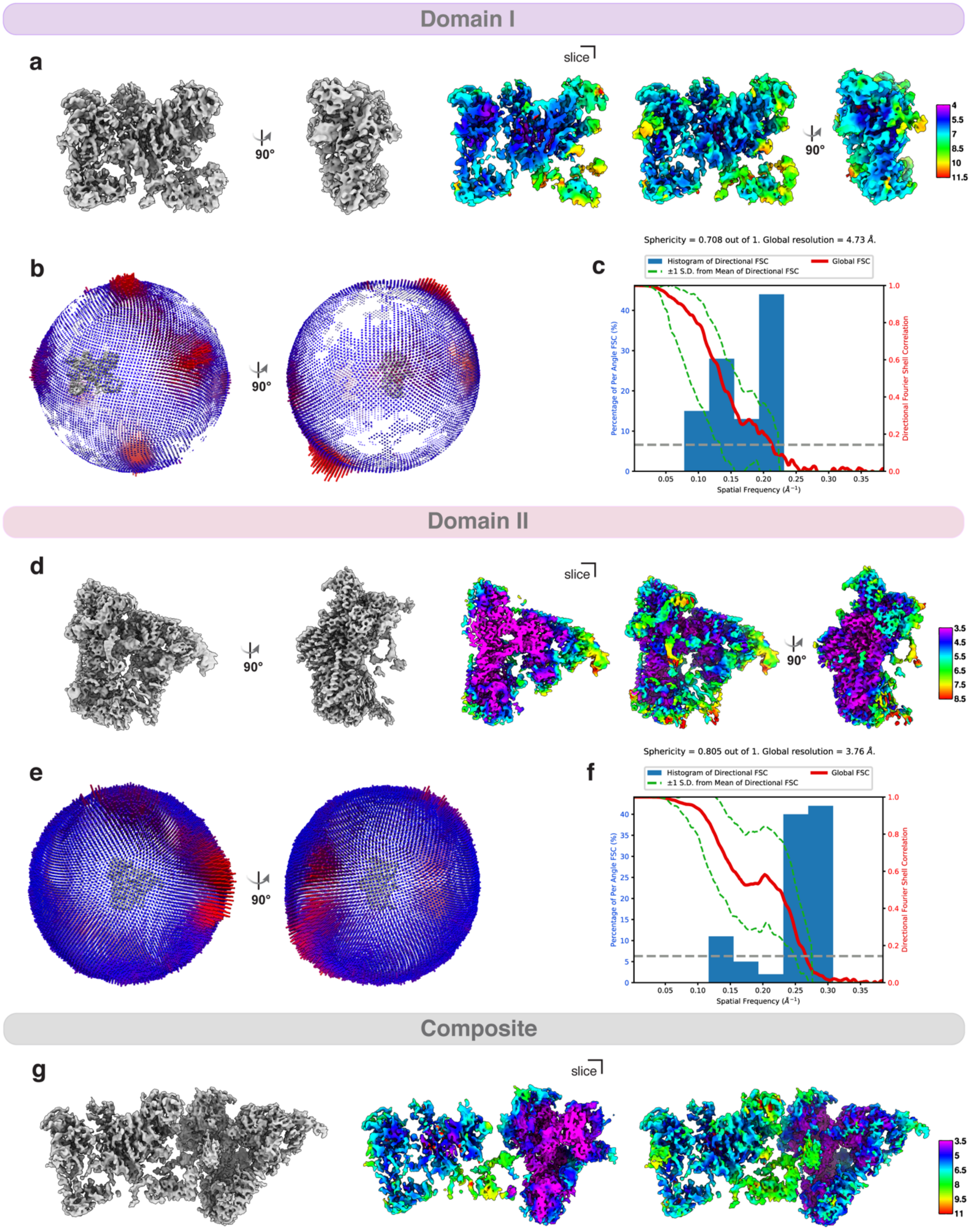

Domain I
a
slice
90°
90°
4
5.5
7
8.5
10
11.5
b
90°
c
Sphericity = 0.708 out of 1. Global resolution = 4.73 Å.
Histogram of Directional FSC
±1 S.D. from Mean of Directional FSC
Global FSC
Percentage of Per Angle FSC (%)
Directional Fourier Shell Correlation
Spatial Frequency (Å⁻¹)
Domain II
d
slice
90°
90°
3.5
4.5
5.5
6.5
7.5
8.5
e
90°
f
Sphericity = 0.805 out of 1. Global resolution = 3.76 Å.
Histogram of Directional FSC
±1 S.D. from Mean of Directional FSC
Global FSC
Percentage of Per Angle FSC (%)
Directional Fourier Shell Correlation
Spatial Frequency (Å⁻¹)
Composite
g
slice
3.5
5
6.5
8
9.5
11

**Extended Data Fig. 4. Local resolution estimation and 3DFSC curves of the Noc1-Noc2 RNP.**

Local resolution estimation displayed in rainbow color on the cryo-EM map of **(a)** domain I, **(d)** domain II, **(g)** composite map shown in 2 different views as calculated by cryoSPARC. Euler angular distribution of the particles from the final refinement of **(b)** domain I, **(e)** domain II. 3DFSC curve for **(c)** domain I and **(f)** domain II reconstructions calculated by cryoSPARC.

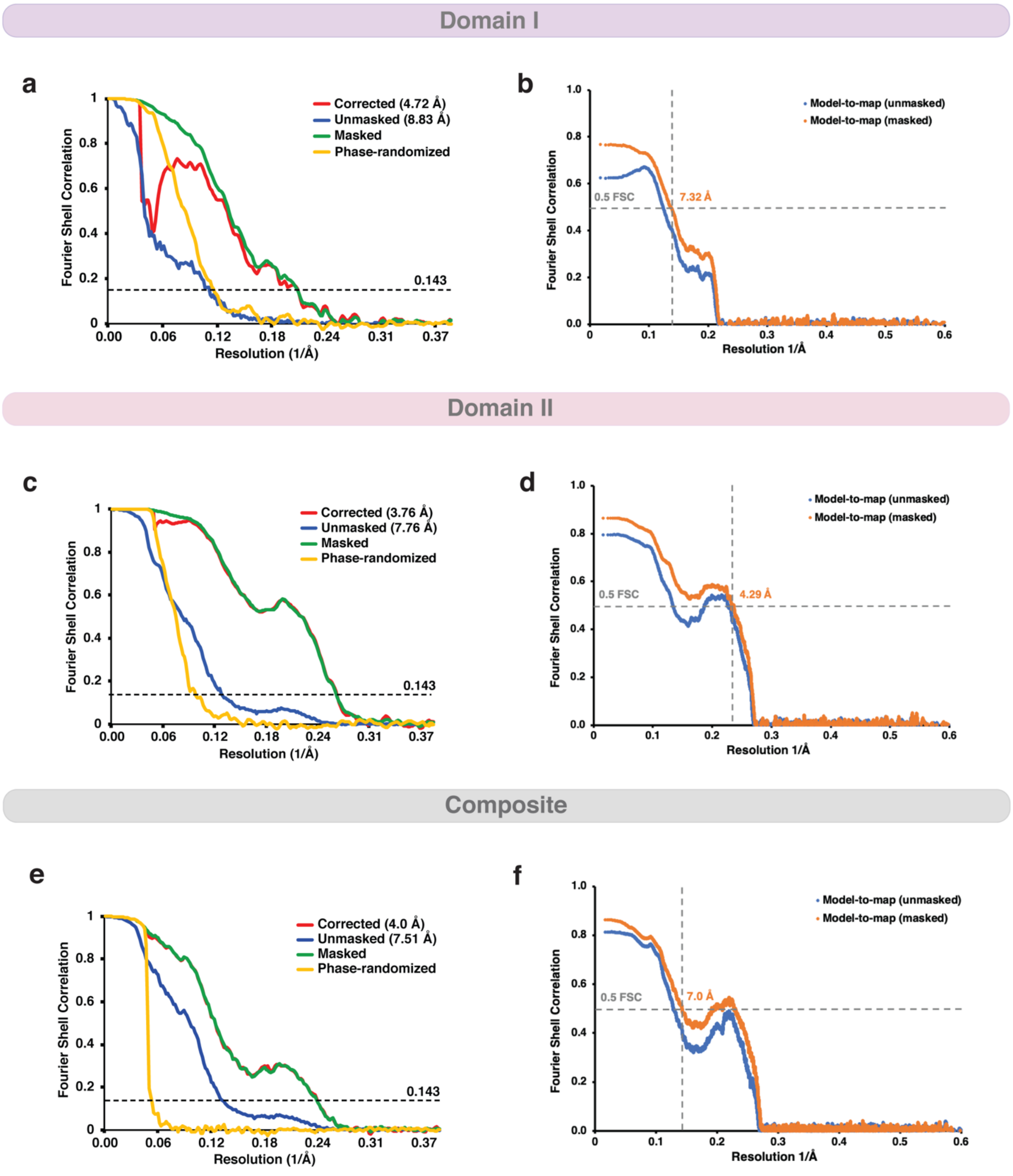


**Extended Data Fig. 5. Solvent-corrected FSC curves and Map-to-Model correlation.**

Solvent-corrected FSC curves for **(a)** domain I, **(c)** domain II and **(e)** composite map as calculated in Relion 3.0. Reported resolutions were determined at FSC=0.143 upon post-processing in Relion 3.0. Map-to-model correlation between **(b)** domain I map and model, **(d)** domain II map and model and **(f)** composite map and model with resolutions reported at FSC=0.5.

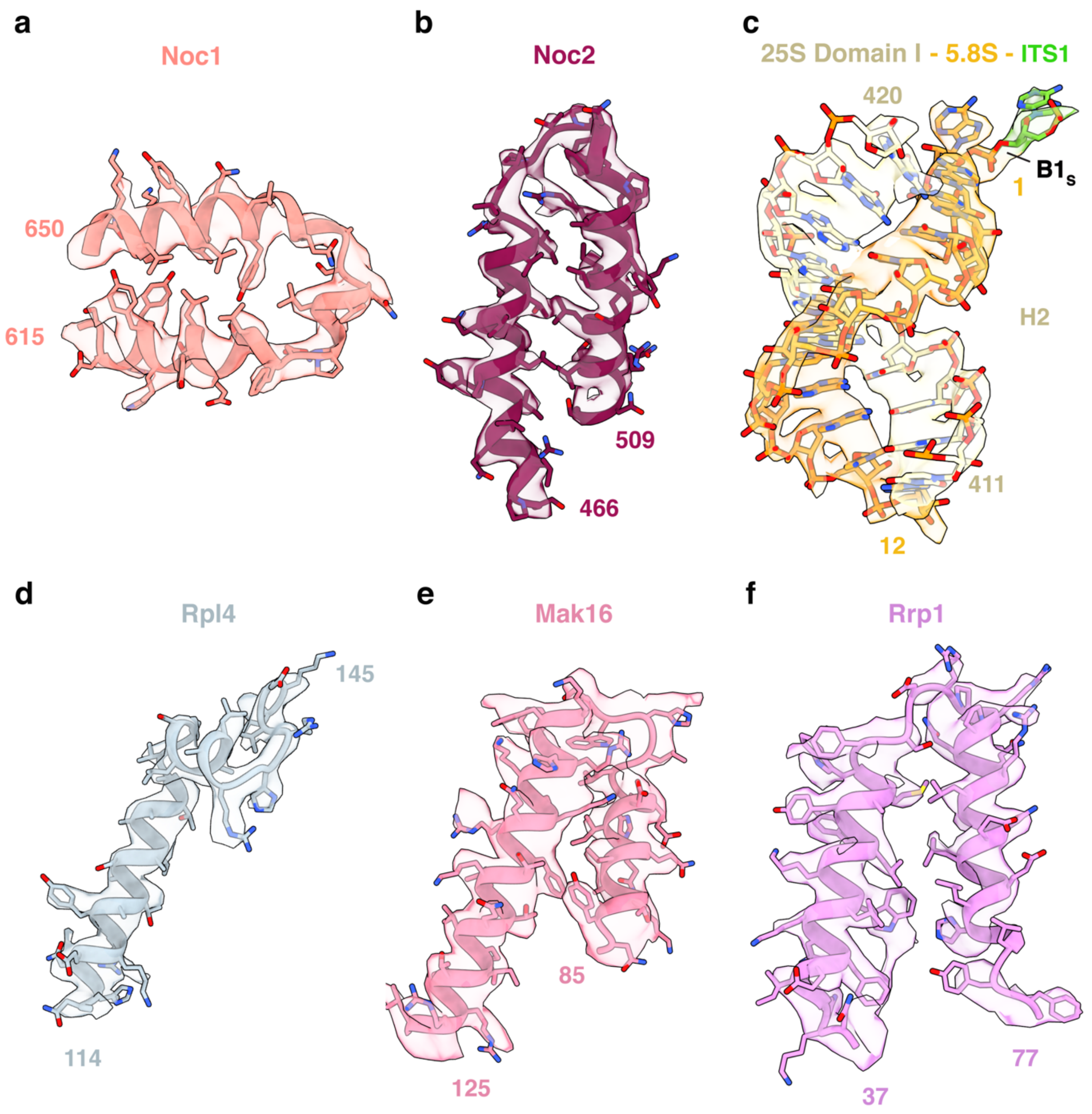


**Extended Data Fig. 6. Representative cryo-EM densities and models for the Noc1-Noc2 RNP.**

**(a-f)** Selections of representative densities of RNA and proteins of the Noc1-Noc2 RNP comprised of assembly factors (a,b,e,f), 25S and 5.8S rRNA from helix 2 (c), and ribosomal proteins (d). Densities are illustrated as continuous transparent volumes and models are shown as ribbons and sticks.

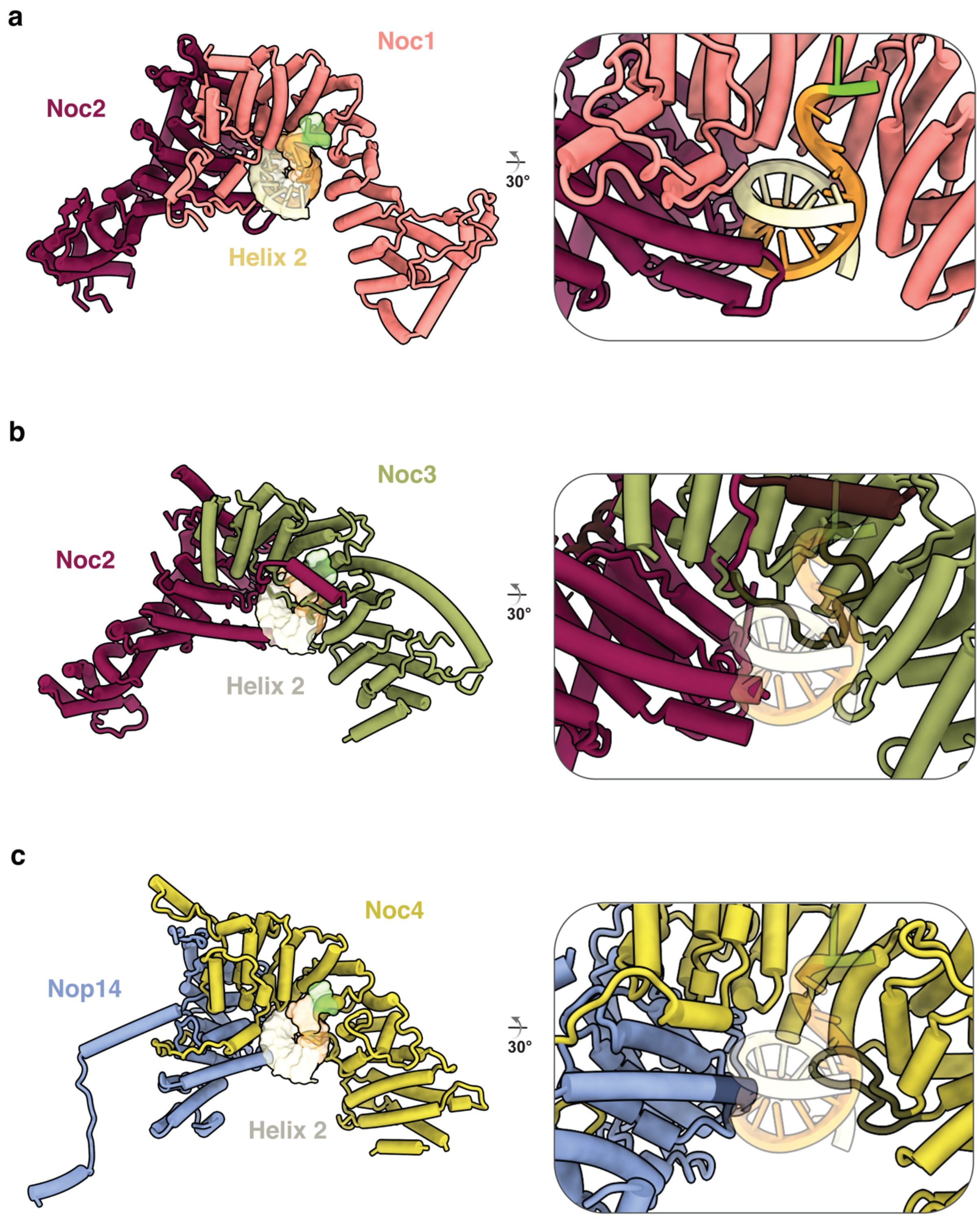


**Extended Data Fig. 7. Comparison of Noc1-Noc2, Noc2-Noc3 and Noc4-Nop14 heterodimers.**

The evolutionarily related helical repeat heterodimers involved in ribosome biogenesis reveal similar structural composition. **(a)** Noc1-Noc2 heterodimer structure has the unique ability to bind to and chaperone an RNA helix structure (helix 2); zoomed inset shows a rotated view. **(b, c)** In comparison, the Noc2-Noc3 heterodimer and the Noc4-Nop14 heterodimer are both unable to bind RNA in a similar fashion owing to steric hinderance from loops in the central cavity, highlighted with darker color.

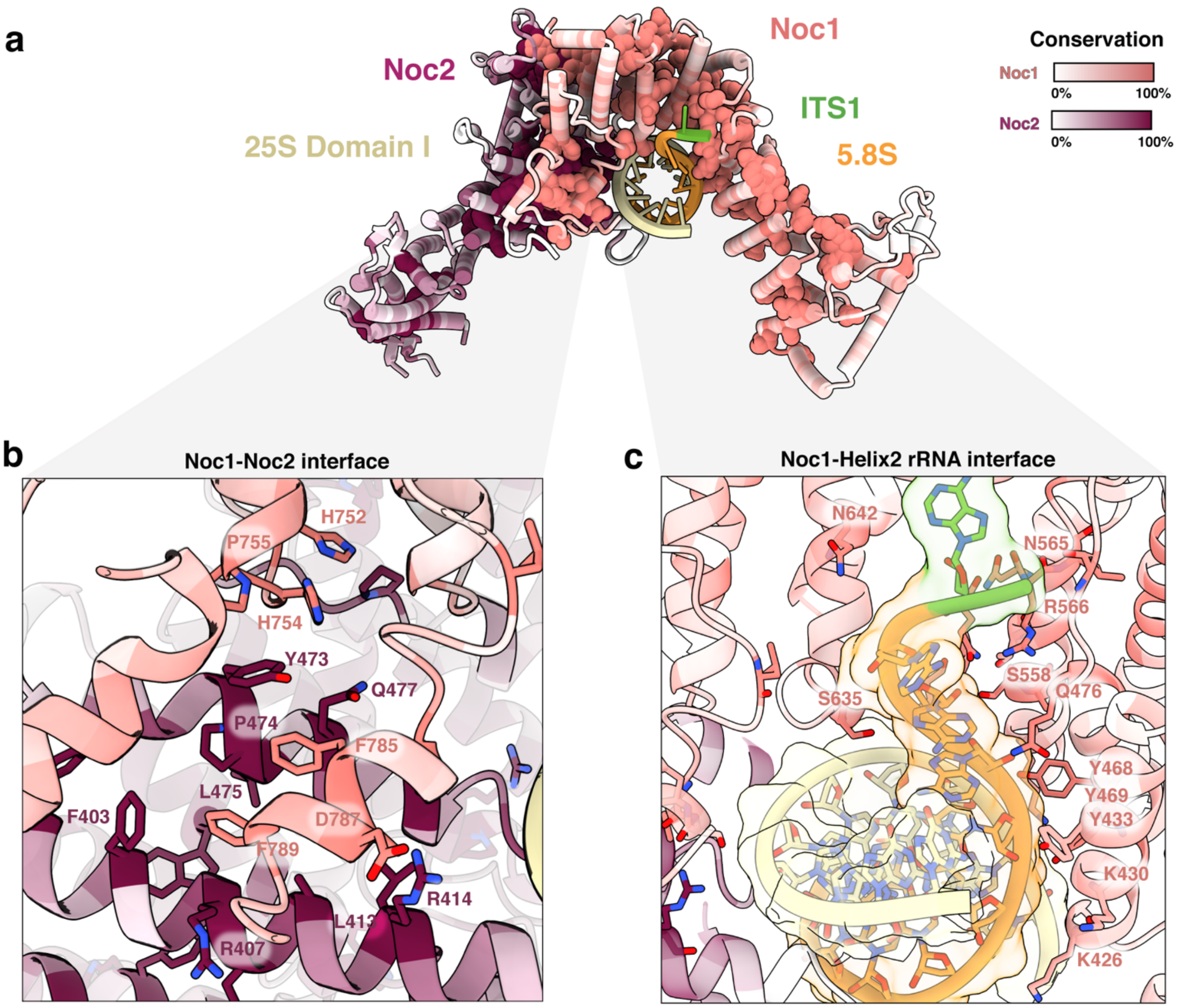


**Extended Data Fig. 8. Sequence conservation highlights critical interfaces of Noc1 and Noc2.**

**(a)** A top view down the central cavity of the Noc1-Noc2 heterodimer encapsulating helix 2 of the large subunit rRNA. The most conserved residues of the heterodimer are depicted in spheres, with sequence conservation colored from coral (100% conserved) to white (0% conserved) for Noc1, and maroon (100% conserved) to white (0% conserved) for Noc2. Conservation was calculated by the ConSurf webserver [37], with sequence alignments generated by Clustal Omega [38] from 12 model organisms. **(b)** The Noc1-Noc2 interface highlights clusters of conserved residues that allow for a tight interaction between the two proteins, with conserved residues depicted in sticks. **(c)** The Noc1-RNA interface reveals a set of conserved residues that line the central channel occupied by the RNA of helix 2, depicted in sticks.

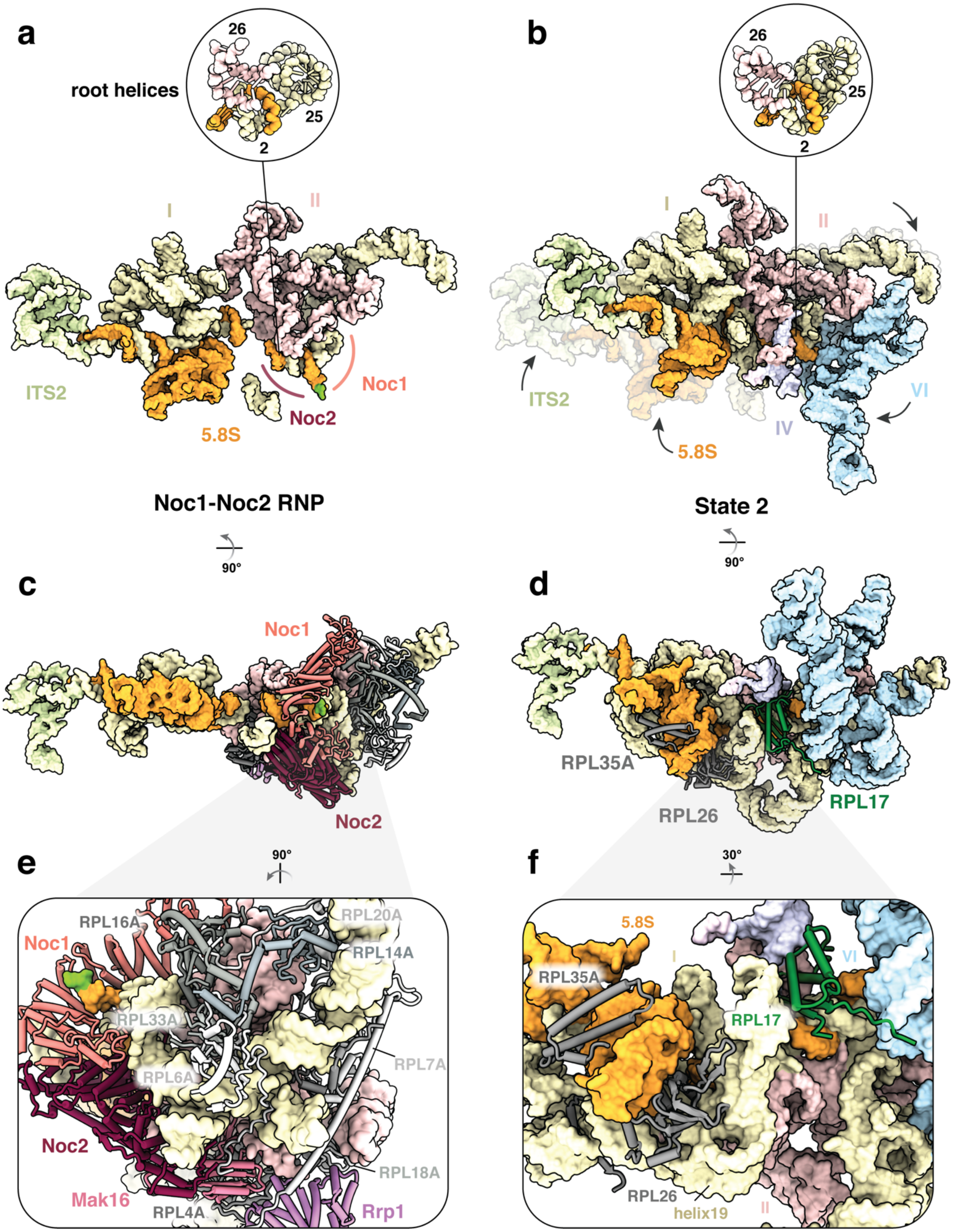
a
root helices
26
25
2
I
II
ITS2
5.8S
Noc2
Noc1
Noc1-Noc2 RNP
b
26
25
2
I
II
ITS2
5.8S
IV
VI
State 2
90°
90°
c
Noc1
Noc2
d
RPL35A
RPL26
RPL17
e
90°
RPL16A
Noc1
RPL20A
RPL14A
RPL33A
RPL7A
RPL6A
Noc2
RPL18A
Mak16
RPL4A
Rrp1
f
30°
5.8S
I
VI
RPL35A
RPL17
RPL26
helix19
II

**Extended Data Figure 9. Analysis of pre-rRNA folding during co- and post-transcriptional ribosome assembly.**

**(a, b)** Comparative view of color-coded rRNA domains within (a) the Noc1-Noc2 RNP and (b) State 2 (PDB 6C0F) with insets highlighting labelled root helices. **(c)** Rotated view with respect to (a). Ribosome assembly factors and ribosomal proteins that stabilize the domain II module are shown as cartoons. **(d)** Rotated view with respect to (b). Ribosomal proteins present at interfaces between rRNA domains are shown as cartoons. (**e**) Zoomed view showing ribosomal proteins stabilizing domains I and II within the Noc1-Noc2 RNP. (**f**) Zoomed view of State 2 (PDB 6C0F) showing ribosomal proteins that stabilize the interface between domains I, II and 5.8S rRNA. Rpl17 is highlighted in green.

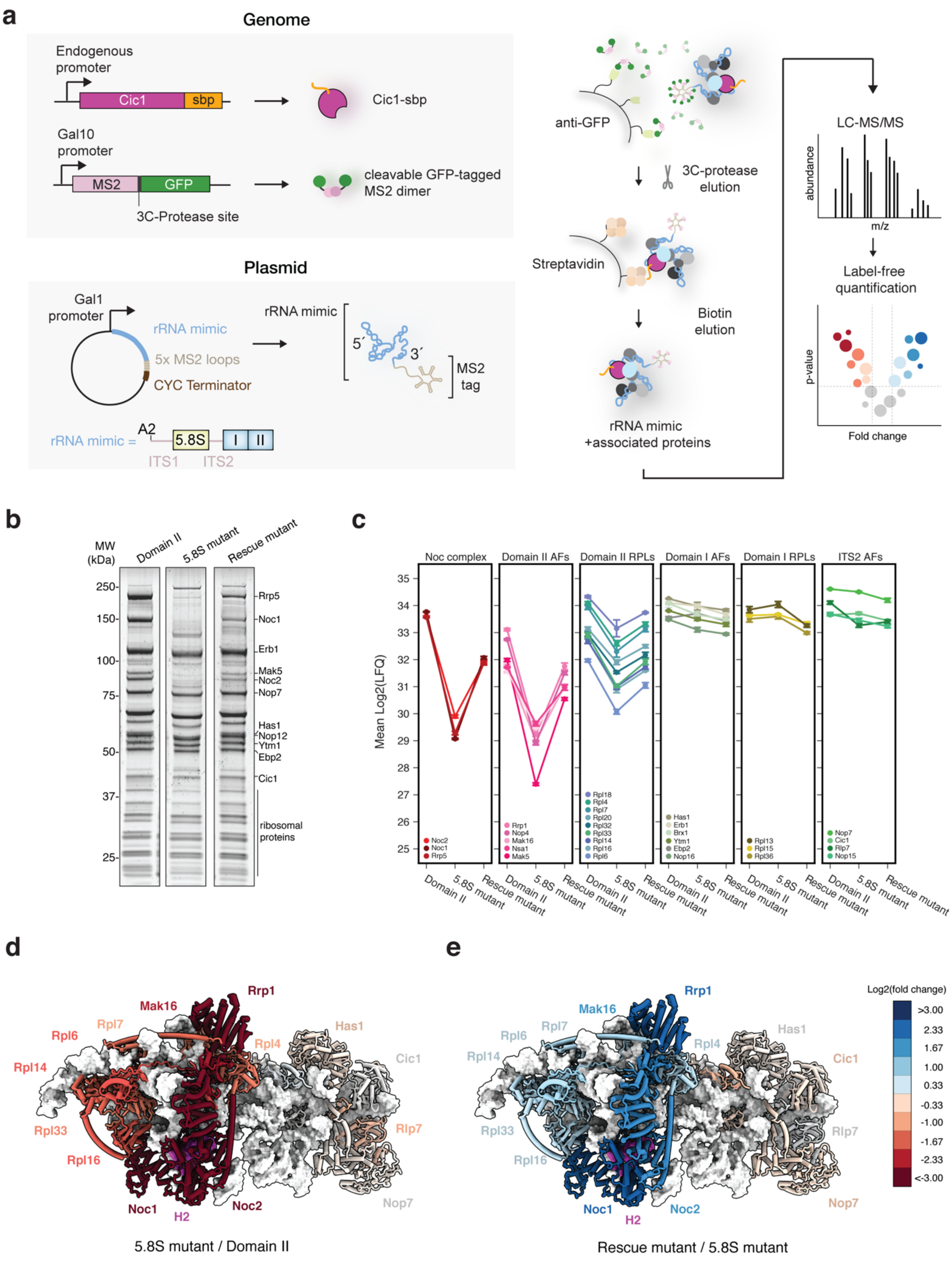
a
Genome
Endogenous promoter
Cic1
sbp
Cic1-sbp
Gal10 promoter
MS2
GFP
cleavable GFP-tagged MS2 dimer
3C-Protease site
Plasmid
Gal1 promoter
rRNA mimic
5x MS2 loops
CYC Terminator
rRNA mimic
MS2 tag
rRNA mimic =
A2
5.8S
I
II
ITS1
ITS2
anti-GFP
3C-protease elution
Streptavidin
Biotin elution
rRNA mimic +associated proteins
LC-MS/MS
abundance
m/z
Label-free quantification
p-value
Fold change
b
MW (kDa)
Domain II
5.8S mutant
Rescue mutant
250
150
100
75
50
37
25
Rrp5
Noc1
Erb1
Mak5
Noc2
Nop7
Has1
Nop12
Ytm1
Ebp2
Cic1
ribosomal proteins
c
Noc complex
Domain II AFs
Domain II RPLs
Domain I AFs
Domain I RPLs
ITS2 AFs
Mean Log2(LFQ)
d
Rrp1
Mak16
Rpl7
Rpl6
Rpl4
Has1
Rpl14
Cic1
Rpl33
Rlp7
Rpl16
Noc1
H2
Noc2
Nop7
5.8S mutant / Domain II
e
Log2(fold change)
Rescue mutant / 5.8S mutant

**Extended Data Figure 10. Disruption of helix 2 prevents assembly of domain II of the 25S rRNA.**

**(a)** Schematic depiction of two-step purification of Domain II mimics followed by mass-spectrometry analysis. Purification of the mimics has been performed similarly to purification of Noc1 RNP except endogenous Cic1 has been C-terminally tagged with streptavidin-biding peptide (sbp). Following purification, samples were analyzed by liquid chromatography-tandem mass spectrometry (LC-MS/MS) and obtained data normalized using label-free quantification. **(b)** SDS-PAGE analysis of purified Domain II rRNA mimics. **(c)** Changes in mean protein abundance, expressed as LFQ values, between purified rRNA mimics: Domain II, 5.8S mutant disrupting helix 2, and rescue mutant restoring helix formation. Error bars represent SD of 3 replicates. **(d, e)** Structure of Noc1-Noc2 RNP colored accordingly to log2 fold change in protein abundance between **(d)** 5.8S mutant and Domain II and **(e)** rescue mutant and 5.8S mutant. Proteins are depicted as cartoons and rRNA as white surface. Helix 2 (H2) is shown in magenta.

### Supplementary Table 1.

| | *Noc1-Noc2 RNP* | *Noc2-Noc3 RNP* |
|---|---|---|
| Data collection and processing | | |
| Magnification | 22,500X | 22,500X |
| Voltage (kV) | 300 | 300 |
| Pixel size (Å) | 1.3 | 1.3 |
| Electron exposure (e- / Å$^2$) | 37.9 | 37.9 |
| Defocus range (um) | -1.0 to -3.0 | -1.5 to -3.0 |
| Number of frames (no.) | 32 | 32 |
| Symmetry imposed | C1 | C1 |
| Initial particle images | 977,232 | 669,811 |
| Final particle images | 158,915 | 75,358 |
| Resolution (Å) | | |
| Global (Å, at FSC 0.143) | 4.0 | 3.69 |
| Local (range, Å, at FSC 0.5) | 2.8-11.5 | 2.8-8.5 |
| Map sharpening B-Factor (Å$^2$) | -44.7 | -28.4 |
| Refinement | ***Phenix_real_space_refine 1.19*** | - |
| Composition (#) | | |
| Chains | 30 | |
| Atoms | 55,125 | |
| Residues | | |
| Protein – | 5158 | |
| Nucleic Acid – | 968 | |
| Water | 0 | |
| Ligands | Zn – 1, Mg – 33 | |
| Bonds (RMSD) | | |
| Length (Å) (# > 4$\sigma$) | 0.004 (0) | |
| Angles (°) (# > 4$\sigma$) | 0.927 (5) | |
| MolProbity score | 1.44 | |
| Clash score | 3.96 | |
| Ramachandran plot (%) | | |
| Outliers | 0.10 | |
| Allowed | 3.69 | |
| Favored | 96.21 | |
| Rotamer outliers (%) | 0 | |
| C$\beta$ outliers (%) | 0 | |
| Peptide plane (%) | | |
| Cis proline/general | 2.3/0.1 | |
| Twisted proline/general | 0.0/0.1 | |
| C$\alpha$BLAM outliers (%) | 2.51 | |
| ADP (B-factors) min/max/mean | | |
| Protein | 83.71/491.41/148.87 | |
| Nucleotide | 88.73/863.92/217.36 | |
| Ligand | 51.94/257.32/115.29 | |
| Water | - | |
| Data | | |
| Supplied Resolution (Å) | 3.9 | |
| Resolution Estimates (Å) | Masked | |
| d FSC (half maps; 0.143) | 4.0 | |
| d 99 (full) | 4.1 | |
| d model | 3.7 | |
| d FSC model (0, 0.143, 0.5) | 3.6/3.7/7.0 | |
| Map min/max/mean | -21.76, 50.64, 0.88 | |
| Model vs. Data | | |
| CC (mask) | 0.64 | |
| CC (box) | 0.73 | |
| CC (peaks) | 0.51 | |
| CC (volume) | 0.64 | |
| RNA Validation | | |
| Average suiteness (%) | 53.2 | |
| Good sugar puckers (%) | 98.55 | |

**Supplementary Table 1. Cryo-EM data collection parameters and refinement and validation statistics.**

### Supplementary Table 2.

| Subgroup | Chain ID | SegID | Molecule name | Total residues or bases | Modelled (residue range) | Initial PDB template |
|---|---|---|---|---|---|---|
| RNA | 1 | L1 | 25S | 3,396 | atomic (1-35, 51-131, 136-168, 251-280, 286-337, 378-393, 407-446, 489-705, 721-751, 778-801,941-952, 1161-1195, 1310-1443) | 6C0F |
| | 2 | L2 | 5.8S | 158 | atomic (1-16, 24-96, 111-158) | 6C0F |
| | 3 | L3 | ITS2 | 232 | atomic (1-67, 213-232) | 6C0F |
| Ribosomal proteins | C | LC | Rpl4A_uL4 | 362 | atomic (2-46, 110-347) | 6C0F |
| | E | LE | Rpl6A_eL6 | 176 | atomic (7-107, 135-176) | 6C0F |
| | e | SE | Rpl32_eL32 | 130 | atomic (52-129) | 6C0F |
| | F | LF | Rpl7A_uL30 | 244 | atomic (3-244) | 6C0F |
| | f | SF | Rpl33A_eL33 | 107 | atomic (4-18, 24-52, 64-107) | 6C0F |
| | G | LG | Rpl8A_eL8 | 256 | side-chain trimmed (53-239) | 6C0F |
| | i | SI | Rpl36A_eL36 | 100 | side-chain trimmed (25-100) | 6C0F |
| | L | LL | Rpl13A_eL13 | 199 | side-chain trimmed (52-127) | 6C0F |
| | M | LM | Rpl14A_eL14 | 138 | atomic (11-125) | 6C0F |
| | N | LN | Rpl15A_eL15 | 204 | side-chain trimmed (2-68, 96-176) | 6C0F |
| | O | LO | Rpl16A_uL13 | 199 | atomic (3-58, 73-199) | 6C0F |
| | Q | LQ | Rpl18A_eL18 | 186 | atomic (15-146) | 6C0F |
| | S | LS | Rpl20A_eL20 | 172 | atomic (2-167) | 6C0F |
| Assembly factors | K | LK | Cic1 | 376 | side-chain trimmed (31-51, 71-303) | 6C0F |
| | n | SN | Nop7 | 605 | side-chain trimmed (13-43, 61-267, 351-460) | 6C0F |
| | o | SO | Nop15 | 220 | side-chain trimmed (88-220) | 6C0F |
| | t | ST | Rlp7 | 322 | side-chain trimmed (54-105, 127-214, 227-322) | 6C0F |
| | s | SS | Erb1 | 807 | side-chain trimmed (239-310, 331-395) | 6C0F |
| | D | LD | Mak16 | 306 | atomic (2-131, Zn-308) | 6C0F |
| | p | SP | Has1 | 505 | side-chain trimmed (42-252, 264-489) | 6C0F |
| | b | SB | Brx1 | 291 | side-chain trimmed (31-194, 211-263) | 6C0F |
| | m | SM | Ebp2 | 427 | side-chain trimmed (196-257) | 6C0F |
| | z | SZ | Rrp1 | 278 | atomic (1-186, 197-253) | 6C0F |
| | 5 | L5 | Noc1 | 1025 | atomic (213-503, 550-701, 744-795, 832-866) | de novo/Alphafold |
| | 6 | L6 | Noc2 | 710 | atomic (175-215, 220-692) | de novo/Alphafold |
| Metal ions | Y | LY | Magnesium | 33 | | 6C0F |

**Supplementary Table 2. Molecular models of the Noc1-Noc2 RNP.**

Individual protein and RNA chains are listed with their initial PDB template (if not built *de novo*), and clustered into categories of RNA, ribosomal proteins and assembly factors.
